# Extended amygdala orchestrates social motivation in socially isolated mice

**DOI:** 10.64898/2026.07.23.739873

**Authors:** Jordan Grammer, Amanda Dryer, Caitlin Tweed, Michael Conoscenti, Moriel Zelikowsky

## Abstract

Juveniles deprived of social contact for an extended period of time demonstrate a broad range of adverse effects on behavior, including increased social aversion. Employing our recently developed integrative assay to assess social aversion, we discovered that social isolation is sufficient to induce decreases in social motivation and increases in social fear and hesitancy. Unsupervised behavioral analyses revealed that social aversion in isolated mice is largely driven by an inflexible, anxiety-like state marked by heightened social vigilance. Using intersectional, projection-specific neural circuit perturbations and in vivo imaging, we discovered that isolation-induced decreases in social motivation are governed by an underexplored projection from the anterodorsal bed nucleus of the stria terminalis (adBNST) to the nucleus accumbens (NAc). These findings illustrate the complexity with which juvenile isolation serves to negatively impact social behavior and identify a novel circuit through which animals integrate information about social state to orchestrate changes in social motivation.

## INTRODUCTION

Sociability is vital across many species^1–7^. In contrast, social aversion - a debilitating state marked by heightened anxiety, apprehension, and diminished motivation toward social contact^8–11^ - is a hallmark characteristic of many neuropsychiatric disorders such as social anxiety, autism spectrum disorder, depression, schizophrenia, and avoidant personailty^12,13^. Long-term disruptions in socialization via prolonged isolation are thought to induce a state of social aversion, including critically impacting social motivation^2,6,14–17^, social interactions^18,19^, and fear in social contexts^20^. Although each of these behavioral domains has been studied individually, it has been challenging to assess social aversion wholistically^8,10,21^, as no single assay has previously captured this state and all of its components collectively.

To address this gap, we recently developed Selective Access to Unrestricted Social Interaction (SAUSI), a single behavioral assay that allows for social choice, free social interaction, and hesitancy measurements in mice^8^. Using this assay, we determined that chronic social isolation in adolescents induces a state of social aversion, characterized by behavioral changes spanning social motivation, fear, and hesitancy^8^. However, the neural mechanisms which enable animals to integrate information about their social state to drive changes in social aversion behaviors remains poorly understood.

The dorsal bed nucleus of the stria terminalis (dBNST) has long been implicated in sustained fear and anxiety-related behaviors, especially in response to stress^6,22–35^. More recent work has expanded this role, demonstrating the dBNST to be socially-sensitive: involved in social vigilance, prosocial behaviors, aggression, sexual behavior, and parenting^36–38^. Moreover, we previously found that this region is responsive to the social state of an animal^6^. This places the dBNST in an ideal position to integrate information related to the social state of an animal in order to drive subsequent changes in behavior. Importantly, distinct dBNST subregions, cell populations, and projections are thought to have heterogenous, or even opposing impacts on behavior^30,33,39^. Specifically, the anterodorsal portion of the dBNST (adBNST) (excluding the oval nucleus), has been shown to have an anxiolytic effect on behavior, while the oval nucleus of the dBNST has been shown to have an anxiogenic effect on behavior^26,30,33,39^. These opposing effects appear to be further shaped by downstream projections^30,40,41^.

Downstream of the dBNST, the nucleus accumbens (NAc) has been repeatedly implicated in both social approach and avoidance behaviors^42–47^. Prolonged social isolation inhibits activity of the NAc^19,48^, and thus, this region is also a prime candidate for mediating social aversion behaviors produced by isolation. Interestingly, the dBNST is thought to send projections to the NAc^49–51^, and stimulating GABAergic, somatostatin^+^ dBNST terminals in the NAc was shown to produce anxiolytic effects^49^. However, this projection – and its broader function – remains otherwise unexplored.

Using SAUSI combined with deep-learning tools^52^ to assess behavior, we found that social isolation produces a myriad of aversive behaviors enhanced by social vigilance, and marked by behavioral inflexibility. Subsequent loss-of-function perturbations and in vivo imaging targeting the adBNST→NAc projection found that this pathway encodes and is necessary for social motivation specifically. Collectively, these results expand our understanding of the impacts of social isolation on social behavior and reveal a projection-specific neural pathway that selectively governs distinct components of isolation-induced social aversion.

## RESULTS

### Prolonged social isolation in juveniles induces a state of social aversion

We tested the impact of prolonged social isolation on mouse behavior using SAUSI^8^, our novel assay to test social aversion in mice, combined with manual annotation and machine learning-based analyses of behavior (Figure 1A). Briefly, adolescent mice were group housed or isolated for four weeks and tested using the SAUSI apparatus, which consists of two outer chambers (home chamber and social chamber) connected by a long narrow tunnel.

**Figure 1.**
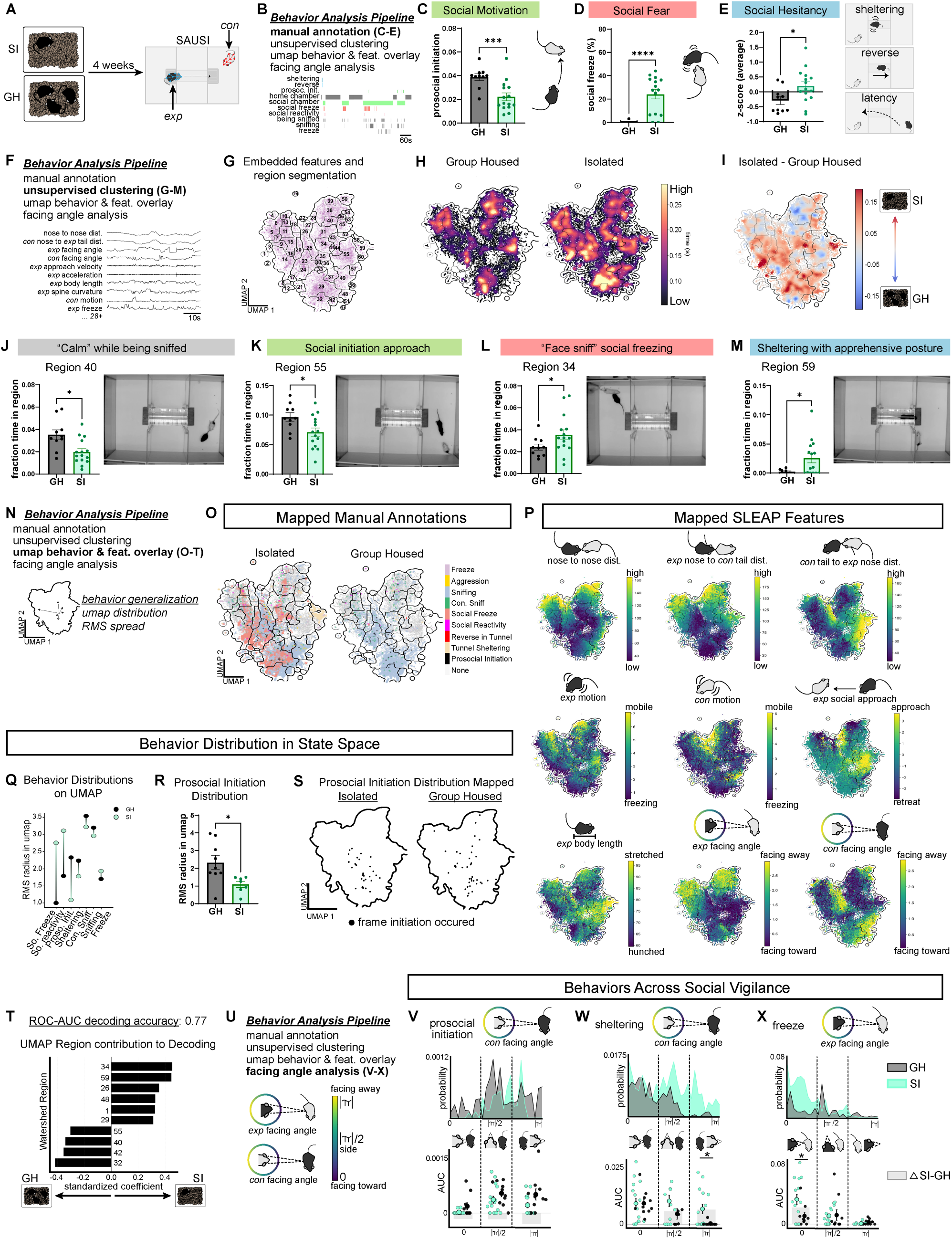
Prolonged social isolation in juveniles induces a state of social aversion. (**A**) Experimental design. (**B**) Behavior analysis pipeline, including an example manually annotated behavior ethogram from one SI mouse. (**C**) Social isolation reduces social motivation as indexed by prosocial initiation (Welch’s t-test, p<.001). (**D**) Social isolation increases social fear as indexed by social freezing (Welch’s t-test, p<.001). (**E**) Social isolation increases social hesitancy as indexed by the z-scored average of social chamber latency, time spent reversing in the tunnel, and time spent sheltering in the tunnel upon first approach (Welch’s t-test, p<.05). (**F**) Behavior analysis pipeline with examples of features calculated from SLEAP postural data. (**G**) “Social” UMAP and segmentation from all mice. (**H**) Time occupancy in UMAP regions by housing condition. (**I**) Change in UMAP occupancy between housing conditions. (**J**) SI mice spend less time occupying region 40 than GH mice (Welch’s t-test, p<.05) (left); still frame from behavior video for region 40 (right). (**K**) SI mice spend less time occupying region 55 than GH mice (Welch’s t-test, p<.05) (left); still frame from behavior video for region 55 (right). (**L**) SI mice spend more time occupying region 34 than GH mice (Welch’s t-test, p<.05) (left); still frame from behavior video for region 34 (right). (**M**) SI mice spend more time occupying region 59 than GH mice (Welch’s t-test, p<.05) (left); still frame from behavior video for region 59 (right). (**N**) Behavior analysis pipeline with diagram for quantifying behavior distribution. (**O**) Manual annotations embedded in UMAP. (**P**) Calculated SLEAP features embedded in UMAP. (**Q**) Averaged distribution of each behavior across UMAP by housing condition. (**R**) Socially isolated mice show tighter distribution of prosocial initiation compared to group housed mice (Welch’s t-test, p<.05). (**S**) Visualization of prosocial initiation UMAP distribution differences across housing conditions. (**T**) Logistic regression model ROC-AUC accuracy & region importance for unsupervised decoding of housing condition. (**U**) Behavior analysis pipeline highlighting facing angle analyses. (**V**) Social isolation shifts prosocial initiation behavior to more probable when the conspecific is facing away (top), but does not reach significance comparing AUC (bottom) (main effect of housing condition (F(1,69) = 7.84, p<.01), main effect facing angle (F(2,69) = 7.85, p<0.001), no effect interaction (F(2,69) = 0.30, p=0.7); FDR corrected post-hoc stars shown on graph. (**W**) Socially isolated mice are more likely to shelter than group housed mice when conspecifics are facing away (shown in distribution, top; AUC, bottom) (main effect of housing condition (F(1,69) = 4.65, p=.03), no main effect facing angle (F(2,69) = 2.77, p=.07), no effect interaction (F(2,69) = 1.21, p=0.3); FDR corrected post-hoc stars shown on graph. (**X**) Socially isolated mice are more likely to freeze than group housed mice when they are facing the conspecific (shown in distribution, top; AUC, bottom) (no main effect of housing condition (F(1,69) = 2.76, p=.1), main effect facing angle (F(2,69) = 11.8, p<.0001), interaction effect (F(2,69) = 4.15, p=0.02); FDR corrected post-hoc stars shown on graph. *n=10 group, housed n=15 isolated for all data in* Figure 1*. All error bars represent mean +/- standard error of the mean. GH: group housed; SI: socially isolated; AUC: area under the curve*.

Prior to testing, experimental mice were habituated to the apparatus and conspecifics were pre-trained to stay in the social chamber, as previously described^8^. During testing experimental and conspecific mice were both placed in the apparatus, and behavior was recorded and analyzed. Using manual annotations of behavior (Figure 1B), we found that social isolation reduced social motivation (prosocial initiation, Figure 1C), increased social fear (social freezing, Figure 1D), and increased social hesitancy behaviors (comprised of sheltering in the tunnel during first approach, reversing in the tunnel during approach, and latency to enter the social chamber, Figure 1E), 3 key hallmarks of social aversion^8,10^.

We then further characterized shifts in behavior state^53^ following social isolation using unsupervised analyses of features extracted from Social LEAP Estimates Animal Poses (SLEAP)^54^ postural tracking (Figure 1F). We calculated 38 features from raw SLEAP data spanning body point distances, facing angle, body length, spine curvature, approach, velocity, acceleration, and motion metrics, all of which were included in our unsupervised pipeline (see STAR METHODS). Features were filtered to include “social” frames (when mice were close together – less than two body lengths in distance). To condense and visualize this high dimensional dataset, we embedded these features into a 2-dimensional UMAP and used watershed segmentation to segregate the map into clustered “regions” using data collapsed across all groups^55–58^ (Figure 1G). We identified how housing condition shapes an animal’s behavior state by quantifying and mapping the time spent occupying space in the UMAP for group housed vs. isolated mice (Figure 1H) and found that social isolation produced significant increases and decreases in the occupancy of numerous clusters (Figure 1I). Specifically, socially isolated mice spent less time occupying region 40, which denoted a “calm” demeanor (i.e., no high velocity movements or freezing, no scrunched or elongated posture, no increased vigilance) while being sniffed (Figure 1J), as well as region 55, which represented social approach (i.e., the action of moving toward the conspecific) (Figure 1K). We also found that socially isolated mice spent more time social freezing while being sniffed near the face (region 34, Figure 1L) and sheltering with an apprehensive, outstretched posture (region 59, Figure 1M).

We performed the same analyses with postural data extracted from across the entire assay (social and non-social frames) and found similar outcomes (Figure S1A-C). Group housed mice showed increased occupancy in “calm” demeanor UMAP space (i.e., no high velocity movements or freezing, no scrunched or elongated posture, no increased vigilance) in the presence of a conspecific (region 12, Figure S1D). Occupancy in social approach from long distances clusters (e.g., region 9) was decreased in socially isolated mice (Figure S1E). Ultimately, these differences in UMAP occupancy highlight the nuanced shifts in behavior state following stress that could not be captured by the human eye alone.

To understand how classified behaviors map onto distinct clusters in UMAP space (Figure 1N), we overlayed manual annotations (Figure 1O) onto our UMAP and found that many behaviors spanned multiple regions, indicating that the UMAP segregates more minute differences in behaviors than observed by eye. When we mapped SLEAP features (Figure 1P) onto our UMAP we were able to visualize the motor and postural contributions to behavioral state. Using Root Mean Square (RMS) analysis to quantify whether manually annotated behaviors were tightly clustered or more dispersed across the UMAP, we found that social fear behaviors were more highly dispersed and social motivation behaviors were more tightly clustered in isolated mice (Figure 1Q). Specifically, prosocial initiation was significantly more clustered in isolated mice (Figure 1R-S), indicating that isolated mice were only willing to approach the conspecific under particular social conditions. Across all frames (Figure S1F-G) we similarly found that prosocial initiation was more tightly clustered in isolated mice (Figure S1J-L). We also analyzed the transition rate between watershed regions and found that overall, socially isolated mice tended to transition between regions less often than group housed mice (Figure S1H-I), suggesting a lack of fluidity in social behavior following isolation.

To test whether time spent occupying space on the UMAP could be used to identify an animal’s social state (housing condition), we used a logistic regression model for decoding. We found that our model was able to decode whether mice were group housed or isolated with 77% accuracy (Figure 1T). The highest contributing features were related to “calm” while being sniffed (regions 40, 42, 32), social approach (region 55), social fear (region 34, “face sniff” social freeze; region 29, “tail sniff” social freeze; and region 48 “tail sniff” and retreat), hesitancy (region 59, apprehensive sheltering), and vigilance (region 26, watching conspecific move from afar) (Figure 1T). The ability to decode whether an animal has been group housed or isolated based solely on data obtained from unsupervised analyses suggests that an animals social state is able to permeate across numerous behaviors, postures, and subtle movements.

Lastly, to determine if social vigilance interacts with social isolation to influence behavior, we measured whether the experimental mouse watched or was being watched by the conspecific during aversive behavior using facing angle analyses (Figure 1U). We found that when prosocial initiation occurred in isolated animals, it did so when the conspecific was facing away (Figure 1V). We also found that while sheltering (an index of social hesitancy) was most likely to occur when the conspecific was facing the experimental mouse regardless of housing condition, isolated mice were more likely to shelter when the conspecific was facing away (Figure 1W). This shows a generalized propensity for isolated mice to hesitate before entering an open social arena, regardless of whether or not they are in the visual plane of a conspecific. Finally, socially isolated mice demonstrated higher freezing behavior specifically when they were facing the conspecific (Figure 1X), suggesting that social isolation not only induces increased social freezing in response to being sniffed (Figure 1D,L), but also increases freezing in the context of heightened social vigilance. These results further demonstrate that social isolation induces a state of reduced sociability and increased behavioral rigidity in mice.

### The adBNST is necessary for social motivation

Because previous work has implicated the anterodorsal subregion of the dBNST in social behavior^36,37^, we tested whether the adBNST plays a role in mediating social aversion behaviors using a virally-mediated chemogenetic approach (AAV2-hSyn-hM4D(Gi)-mCherry) to silence the adBNST during social aversion testing (Figure 2A). Group housed mice were tested for social aversion behaviors on SAUSI under the influence of the DREADDs activating ligand Deschloroclozapine (DCZ, 0.25 mg/kg i.p.) to reversibly inhibit the adBNST with no off-target effects (Figure S2A-D). The same animals were tested again following injection of the saline control (order counterbalanced). We found that adBNST silencing selectively impaired social motivation, but was not sufficient to induce social fear or hesitancy (Figure 2B). This suggests that the adBNST serves to drive social motivation in mice, as silencing this region was sufficient to induce an isolation-like social motivation deficit in group housed mice.

**Figure 2.**
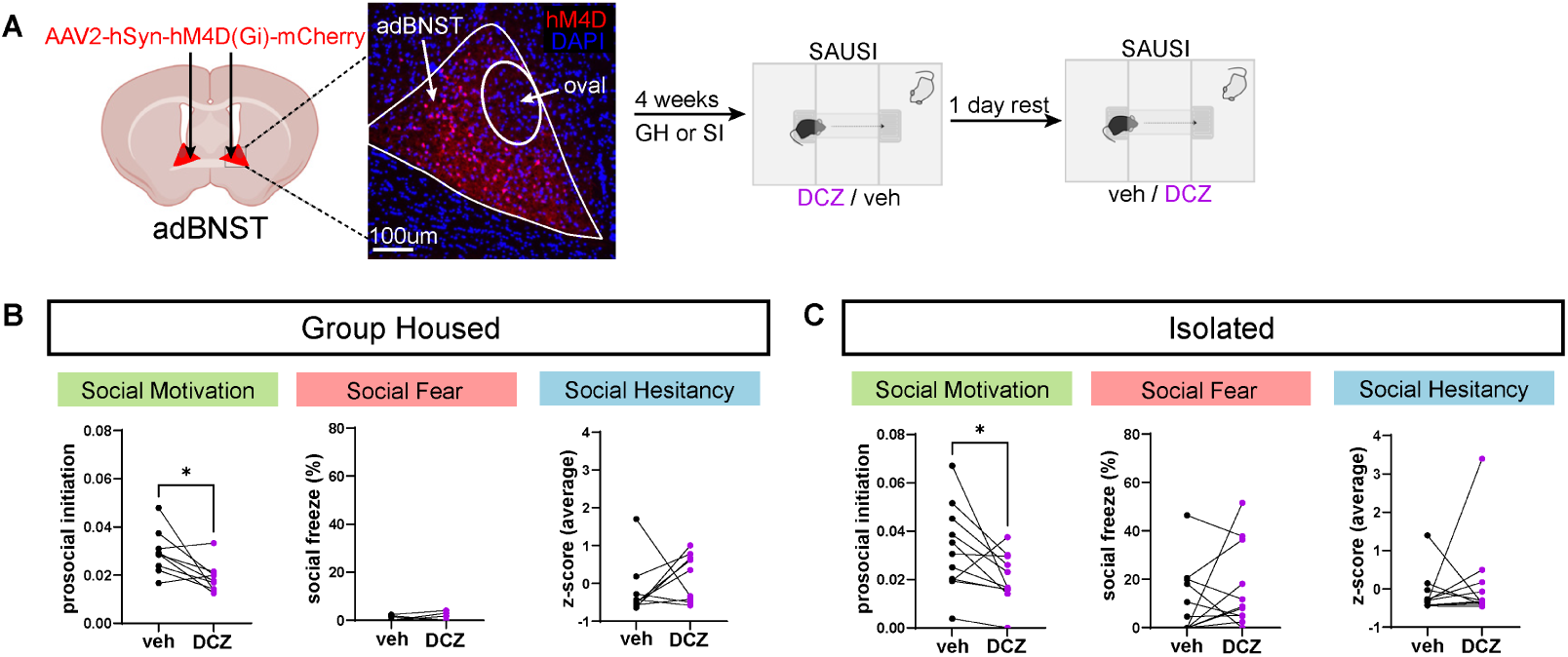
The adBNST is necessary for social motivation. (**A**) Experimental design & viral targeting of adBNST. (**B**) Silencing adBNST in GH mice (n=9) significantly reduces prosocial initiation (p=.02), but does not alter social fear (p=.5) or hesitancy (p=.4). (**C**) Silencing adBNST in SI mice (n=11) significantly reduces prosocial initiation (p=.04), but does not alter social fear (p=.3) or hesitancy (p=.5). *All comparisons paired-samples t-test. All error bars represent mean +/-standard error of the mean. GH: group housed; SI: socially isolated; DCZ: deschloroclozapine; veh: vehicle*.

Consistent with these data, when we chemogenetically silenced the adBNST in isolated mice, we found that social motivation was even further disrupted, with no effects on social fear or hesitancy, again highlighting a role for the adBNST in the regulation of social motivation specifically (Figure 2C). Interestingly, chemogenetic activation of this region in socially isolated mice was not sufficient to recover normal social behavior, including social motivation (Figure S2I-L), suggesting that this region is required for the expression of social motivation but is not sufficient to rescue deficits in social motivation in isolated mice.

Moreover, these data suggest that the behaviors which comprise the state of social aversion – social motivation, fear and hesitancy – may be separably regulated by distinct brain structures or subregions. Indeed, manipulations of the anxiogenic oval nucleus of the dBNST were sufficient to generate a distinct social aversion category – social fear – in group housed mice (Figure S3A-E).

### GABAergic adBNST projections to the NAc are necessary for social motivation

To test whether adBNST control over social motivation is mediated by projections to the NAc, we used a dual virus, intersectional approach to chemogenetically silence neurons in the adBNST that project to the NAc during social aversion testing (Figure 3A). Specifically, we delivered a retrograde Cre-expressing virus (pENN-AAVrg-hSyn-HI-eGFP-Cre-WPRE-SV40) to the NAc and a Cre-dependent DREADDs virus (AAV2-hSyn-DIO-hM4D(Gi)-mCherry) into the adBNST, and after four weeks of group housed or social isolation conditions, tested social aversion behavior with SAUSI. In line with our pan-neuronal adBNST effects, we found that silencing adBNST→NAc reduced social motivation in both group housed (Figure 3B) and isolated mice (Figure 3C), but left other social aversion behaviors intact. These data suggest that the adBNST controls social motivation via projections to the NAc.

**Figure 3.**
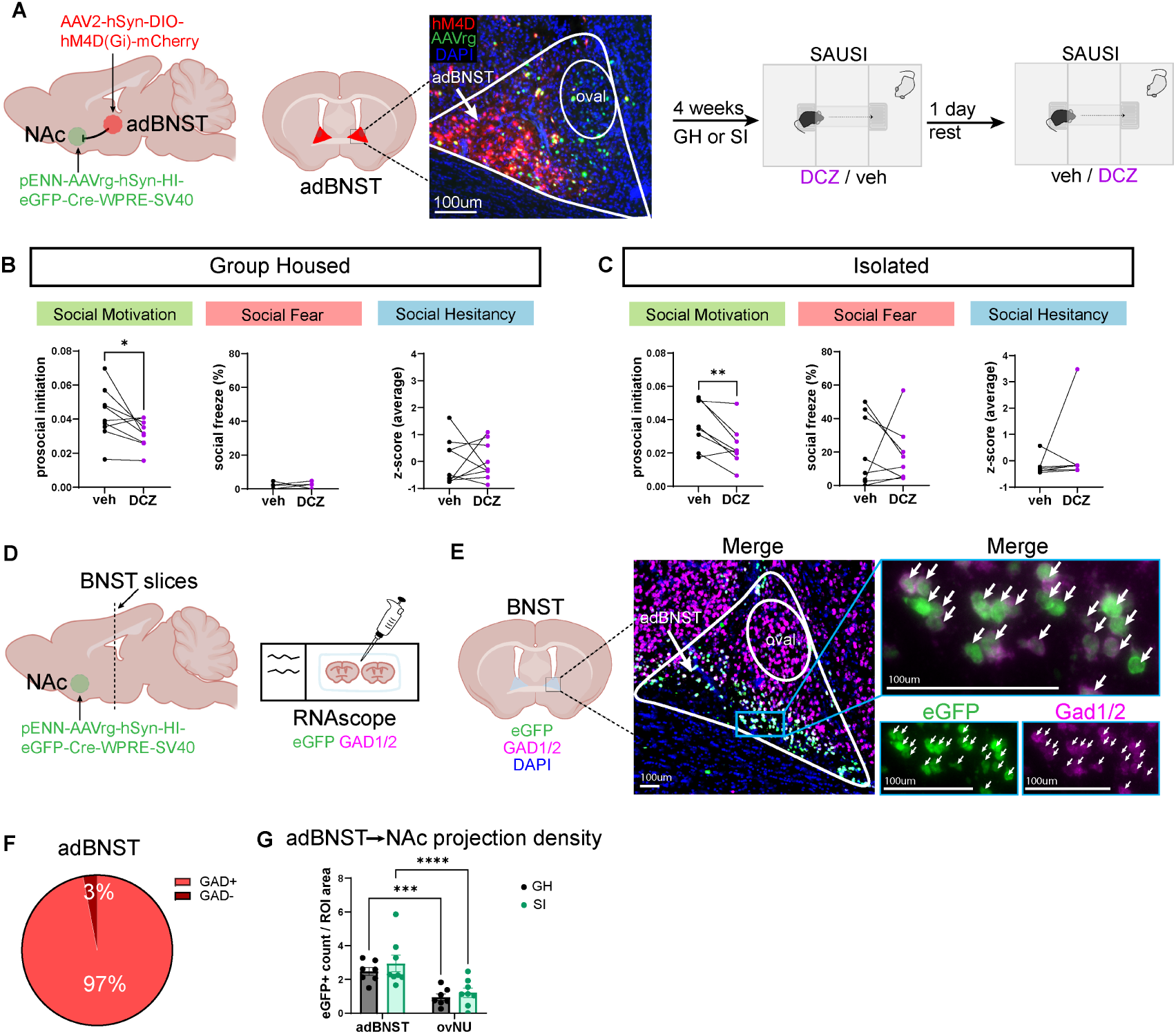
GABAergic adBNST projections to the NAc are necessary for social motivation. (**A**) Experimental design & viral targeting. (**B**) Silencing adBNST→NAc in GH mice (n=9) significantly reduces prosocial initiation (paired t-test, p=.04), but does not alter social fear (paired t-test, p=.9) or hesitancy (paired t-test p=.9). (**C**) Silencing adBNST→NAc in SI mice (n=8) significantly reduces prosocial initiation (paired t-test, p<.01), but does not alter social fear (paired t-test, p=.8) or hesitancy (paired t-test, p=.4). (**D**) Experimental design for adBNST→NAc projection quantification with RNAscope. (**E**) Representative microscopy images of dBNST, with inset images showing colocalization of NAc projectors and GAD1/2. (**F**) Percentage of projections labelled with GAD 1/2. (**G**) dBNST to NAc projections originating from oval nucleus vs anterodorsal subregion by housing condition (no main effect of housing condition (F(1,13) = 0.79, p=.3), main effect subregion (F(1,13) = 51.5, p<.0001), no effect interaction (F(1,13) = 0.22, p=0.6); Holm-Sidak corrected post-hoc stars shown on graph. *Data points represent per mouse averages from 2 hemispheres of a single brain slice each. All error bars represent mean +/- standard error of the mean. GH: group housed; SI: socially isolated; DCZ: deschloroclozapine; veh: vehicle*.

The BNST contains 80-97% GABAergic neurons^24,27,59,60^, and dBNST projections to NAc interneurons are thought to be largely GABAergic^49^ and originate specifically from the ad nucleus of the dBNST^61^. However, the genetic identity of adBNST cells that project to non-specific cell types in the NAc is unknown. To determine the genetic identity of adBNST→NAc projections, we used a Cre-expressing retrograde virus injected into the NAc (pENN-AAVrg-hSyn-HI-eGFP-Cre-WPRE-SV40) to label NAc projectors upstream in the adBNST. Brains were then extracted and processed for GAD1/2 RNA expression using fluorescent in situ hybridization (RNAscope) on adBNST-containing sections (Figure 3D). We found that 97% of projections from the adBNST to NAc were GABAergic (Figure 3E-F). Quantification of projection density revealed that a majority of NAc efferents originate from the anterodorsal subregion rather than the oval nucleus, with no differences between housing condition (Figure 3G). These data confirm that BNST→NAc projectors predominantly originate in GABAergic cells in the adBNST.

### adBNST→NAc ensembles are primarily tuned to social motivation behaviors

Building on our discovery that adBNST→NAc projections are necessary for social motivation behaviors, we tested whether these projectors were uniquely tuned to social motivation behaviors or were tuned to social aversion behaviors more broadly. Mice were injected with a retrograde Cre-expressing mCherry virus (AAV2rg-hSyn-Cre-mCherry-WPRE-hGH-polyA) into the NAc, and a Cre-dependent GCaMP virus (AAV1-hSyn-FLEX-jGCaMP8m-WPRE) into the adBNST. An integrated GRIN Lens (0.6mm x 7.3mm) was implanted over the adBNST to allow us to image calcium activity in adBNST→NAc projectors. To assess ensemble activity during social aversion behaviors, mice were socially isolated following surgery and calcium activity of adBNST→NAc cells was imaged using a miniature microendoscope (nVue) during SAUSI behavior testing (nVision) (Figure 4A). Using peri-event time histograms (PETHs) to measure statistically significant and robust (magnitude >3x baseline) neural activity responses to behavior, we identified cells that were tuned to each social aversion category. Neurons were largely tuned to social motivation behaviors (prosocial initiation (Figure 4B) and social chamber entry (Figure 4C)), with a smaller subset of cells demonstrating selective tuning to social fear behaviors (social freezing (Figure 4D) and social reactivity (Figure 4E)) and social hesitancy behaviors (reversing in the tunnel (Figure S4A-C) and sheltering in the tunnel (Figure S4F-H)). Collectively, adBNST→NAc neurons were primarily tuned to social motivation behaviors – roughly double that of social fear, with minimal overlap between populations (Figure 4F-G).

**Figure 4.**
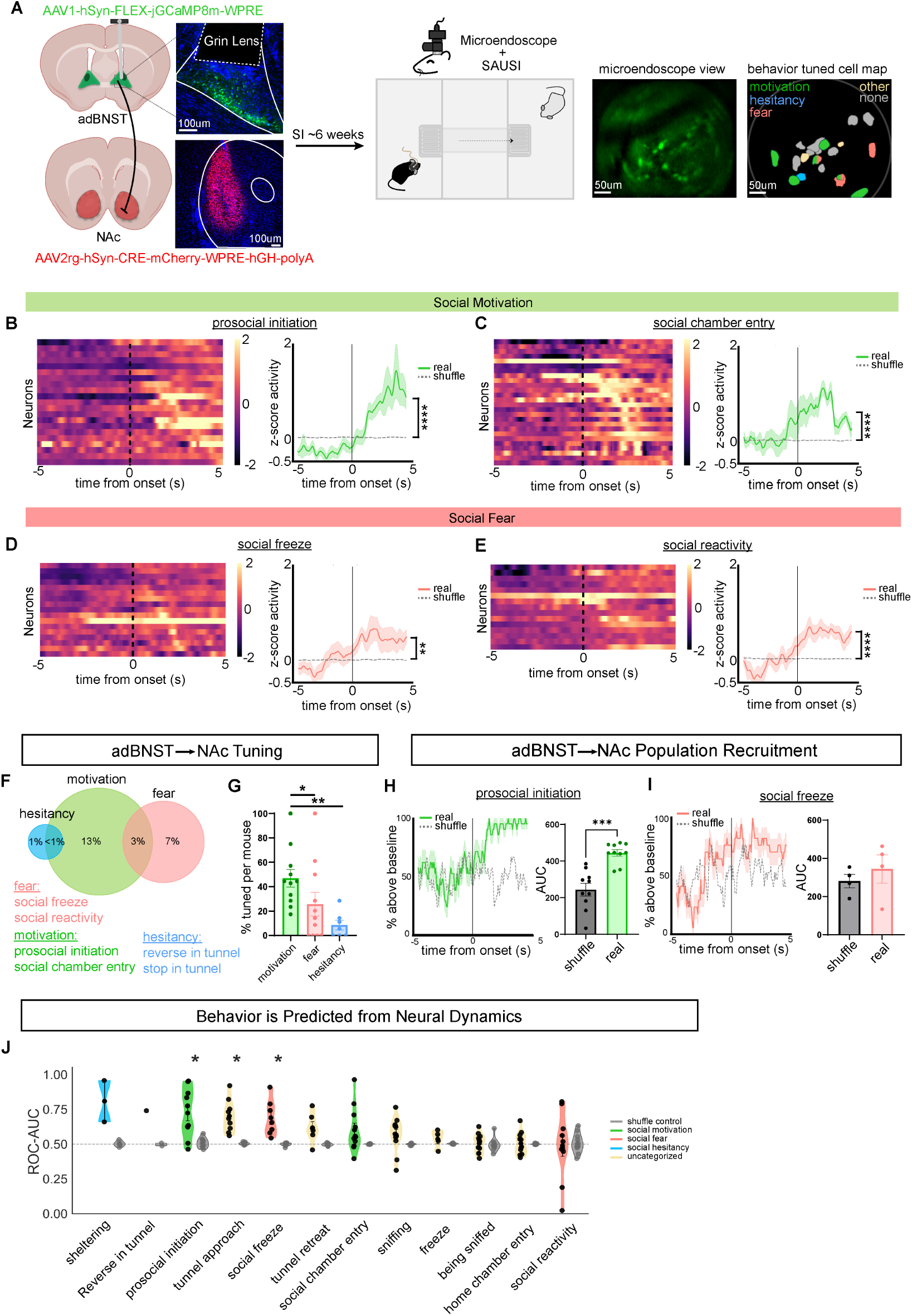
adBNST→NAc ensembles are primarily tuned to social motivation behaviors. (**A**) Experimental design showing viral targeting and lens placement over adBNST. Representative image of focal plane and map depicting cell tuning to behavior categories. (**B**) Heatmap (left) and PETH (right) showing neurons tuned to prosocial initiation (n=21 neurons; Wilcoxon signed-rank test, p<.0001). (**C**) Heatmap (left) and PETH (right) showing neurons tuned to social chamber entry (n=26 neurons; Wilcoxon signed-rank test, p<.0001). (**D**) Heatmap (left) and PETH (right) showing neurons tuned to social freezing (n=17 neurons; Wilcoxon signed-rank test, p<.01). (**E**) Heatmap (left) and PETH (right) showing neurons tuned to social reactivity (n=15 neurons; Wilcoxon signed-rank test, p<.0001). (**F**) Majority of neurons are tuned to social motivation behaviors, with little overlap with other social aversion categories (% of all (tuned and not tuned) neurons in the experiment). (**G**) Significantly more cells are tuned to social motivation compared to social fear and social hesitancy. (% of total tuned neurons per mouse) (Mixed-Effect Model F(1.697, 13.58) = 7.24, p<.01) Holm-Sidak corrected post-hoc comparisons. (**H**) Tuned neurons show coordinated population activity in response to prosocial initiation (left) and significantly increased post-onset response (AUC comparison (right) (n=10 mice; paired t-test, p<.001)). (**I**) Tuned neurons show no coordinated population activity in response to social freezing (left) and no difference in post-onset response (AUC comparison (right) (n=4 mice; paired t-test, p=.4)). (**J**) Lagged logistic regression modeling significantly predicts prosocial initiation (n=10 mice; FDR corrected paired t-test, p=.027), tunnel approach (n=9 mice; FDR corrected paired t-test, p=.012), and social freezing (n=7 mice; FDR corrected paired t-test, p=.028) above chance, with accuracy indexed as ROC-AUC. *n=253 total neurons recorded from n=11 mice. Individual data points represent values per mouse. All error bars represent mean +/- standard error of the mean. SI: socially isolated; AUC: area under the curve; ROC-AUC: receiver operating characteristic area under the curve*.

To determine whether socially tuned cells had coordinated population activity in response to behavior we used PETHs to measure the percent of tuned neurons that were active in response to behavior (activity > average pre-onset activity). We found that prosocial initiation recruited approximately 100% of tuned cells after onset, which was significantly increased compared to a shuffled control (Figure 4H). In contrast, social freezing did not show a significant increase in population recruitment after onset (Figure 4I). Similarly, social hesitancy behaviors did not have distinguishable population recruitment compared to shuffle controls (Figure S4D-E,I-J). This suggests a coordinated population response specifically to social motivation behavior, which provides further evidence for the specificity of our perturbation manipulations (Figure 3).

Lastly, to test whether social motivation can be more easily decoded from neural activity than other behaviors, we applied a lagged logistic regression model to predict whether a given behavior was occurring based on calcium activity. We found three behaviors that were predicted with an accuracy significantly higher than chance, including prosocial initiation (mean ROC-AUC 0.72), tunnel approach (mean ROC-AUC 0.70), and social freezing (mean ROC-AUC 0.69). Tunnel approach was not categorized in the initial social aversion phenotype, but is considered a social motivation behavior in the context of neural activity as it indicates the decision to approach the social chamber^8^. Tunnel approach also showed high tuning and population recruitment (Figure S4K-O). Other behaviors related to changes in social context had pronounced neural responses (S4P-Y), but were not decodable from neural activity. The ability to significantly decode social motivation with the highest accuracy from adBNST→NAc ensemble activity, combined with strong tuning and highly coordinated population activity, indicates that this projection primarily responds to social motivation behaviors.

## DISCUSSION

Using our newly innovated, comprehensive behavioral assay SAUSI, we determined that prolonged social isolation induces a state of social aversion which encompasses decreases in social motivation and increases in social fear and hesitancy. Virally-mediated, intersectional perturbations of the adBNST and its projections to the NAc identified a novel and specialized role for this underexplored GABAergic projection in the control of social motivation. In vivo imaging revealed that largely non-overlapping neural ensembles within adBNST→NAc projectors are tuned to distinct behavioral categories of social aversion, with the majority of cells tuned to social motivation. Collectively, our findings delineate a privileged role for GABAergic adBNST→NAc projectors in the control of adaptive social motivation behaviors.

Our manual annotations identified increased social aversion behaviors following isolation. These results were supported and enriched by machine vision-based postural tracking and unsupervised computational analyses, which revealed additional distinctions in how social isolation changes the behavioral state of socially interacting mice. Importantly, we determined that reduced prosocial initiation in isolated mice could not simply be explained by increases in antisocial behaviors such as aggression or ignoring, but because they showed an overall reduction in social approach, identified by UMAP region 55. Further, not only was social initiation reduced by isolation, but the behavioral contexts (clusters) in which social initiation occurred were significantly constricted, suggesting that isolation not only limits prosocial motivation but also increases the rigidity in which social motivation behaviors are displayed.

We similarly identified additional contexts in which isolation-induced changes in social behavior decreased the flexibility of social interactions. In particular, we found that social freezing was broadly distributed across UMAP clusters, but that it was particularly severe in response to being sniffed near the face (region 34), suggesting that isolated mice perseverate in this behavior across a number of behavioral clusters but that it becomes particularly rigid in certain social circumstances. Likewise, tunnel sheltering appeared across several UMAP clusters, but only region 59 - defined by elongated body posture, freezing, and face-to-face orientation between mice - was enhanced by isolation. This specificity suggests that isolation does not only increase hesitancy in general, but constrains it into a stringent, previously unrecognized behavioral configuration. This behavioral inflexibility was also supported by our finding that socially isolated mice tend to transition between UMAP regions less than group housed mice, consistent with a social anxiety-like phenotype and commonly found in autism spectrum disorder^62,63^. Collectively, these unsupervised analyses reveal that social isolation places mice into a state of social aversion that is made up of highly rigid, inflexible behaviors that persist in particular behavioral contexts.

The dBNST has been extensively investigated for its role in anxiety behaviors and stress^6,22–35^. However, more recently, the anterodorsal subregion of the BNST has gained traction for its involvement in sociability^36^, particularly regarding the time spent engaging with a conspecific^37,64–66^. While the dBNST projects to many brain regions involved in social motivation and reward^51^, direct evidence for the adBNST in motivated behaviors is lacking. Here, we directly demonstrate a role for the adBNST in social motivation that cannot be explained by alterations in anxiety, as our adBNST manipulations had no impact on the expression of social fear behaviors^30^. In recent work, Luo et al. failed to find a role for dBNST hypocretin signaling in the control of social approach^67^, however, these manipulations were inclusive of the oval nucleus. Additionally, Yuan et al. found that inhibition of dBNST cells receiving indirect projections from prefrontal cortex rescued anxiety-like behaviors but did not affect social motivation in isolated mice^68^. These data, in combination with the results obtained here, support the idea that subnuclei and circuits of the dBNST are likely highly specialized with distinct and often disparate functions^30^. Indeed, in contrast to our adBNST findings, our oval BNST manipulations were sufficient to drive social fear behaviors, consistent with a role for the oval nucleus in anxiety-like behavior^26,30,33,39^. These findings suggest that distinct components of social aversion – motivational changes, increased social fear, and increased social hesitancy – may be controlled by dissociable but locally integrated subnuclei of the dBNST and related structures. Future work utilizing SAUSI to detect a potential triple dissociation between distinct subregions of the dBNST and distinct social aversion behaviors is warranted.

While recent studies have examined anterior (both dorsal and ventral) BNST projections to reward circuitry^69–72^, very few have directly interrogated the function of dBNST→NAc specifically. To date, Xiao et al. offers the main investigation of this pathway^49^, demonstrating increased functional connectivity between the dBNST and NAc following stress, and identifying an anxiogenic role for a GABAergic, somatostatin-expressing subpopulation of dBNST terminals synapsing onto NAc parvalbumin (PV) interneurons in traditional anxiety-like assays and social recognition tasks. Although this study also tested sociability using the three-chamber assay, it found no differences upon manipulating this pathway. Critically, Xiao et al. used a genetically-restricted, presynaptic viral strategy (targeting GABAergic and, more specifically, somatostatin-expressing dBNST neurons, with no distinction between oval and adBNST) to isolate a molecularly-defined subpopulation within this circuit. This leaves the function of the broader adBNST→NAc projection unknown. Here, we found that adBNST→NAc specifically regulates social motivation in group housed and isolated mice alike.

Neural populations encode a vast array of information to shape internal state and behavior^53^. We found that adBNST→NAc ensembles are largely tuned to approach and retreat from social environments. Beyond encoding social motivation, these data are consistent with the idea that these projections may also participate in a broader network of social context assessment^36^ — one that tracks transitions in social context, such as the start, entry, and exit of a social encounter. Additionally, we did not find significant overlap in the neural ensembles that were tuned to social motivation vs social fear vs social hesitancy, suggesting these separable populations may be genetically distinct and providing motivation for future studies to determine the genetic identity or signaling mechanisms which underly distinct components of social aversion.

Together, these findings deepen our understanding of social aversion behavior in an adolescent social isolation model and identify a dissociable role for adBNST→NAc in social motivation. By resolving dBNST function with subregion and projection-level specificity, this work adds to a growing picture of how distinct circuits within the dBNST contribute to social behavior, and may help inform future therapeutic targets for neuropsychiatric disorders associated with social aversion.

## RESOURCE AVAILABILITY

### Lead contact

Requests for further information, resources and reagents should be directed to the lead contact, Moriel Zelikowsky.

### Materials availability

All items described here are commercially available, or open source^8^.

### Data and code availability

We have made all analysis code for the figures in this paper available at https://github.com/ajgrammer/grammer-et-al-2026.

## ACKNOWLEDGEMENTS

We thank the University of Utah Machine Shop for working with our design to help build the SAUSI apparatus; Rusul Jabber for technical assistance; Damhyeon Kwak for input on SLEAP analysis; Amy Monasterio for consultation with calcium imaging analysis; and Dr. Thomas Kash for generously providing the CRH-Flp mouse line.

## AUTHOR CONTRIBUTIONS

Conceptualization, J.G. and M.Z.; investigation, J.G., A.D., C.T., M.C.; writing J.G. and M.Z.; funding, J.G. and M.Z.; resources, M.Z.; supervision, M.Z.

## FUNDING

This work was supported by the National Science Foundation Graduate Research Fellowship Program (JG), University of Utah Graduate Research Fellowship (JG), a travel scholarship to the *Short Course on the Application of Machine learning for Automated Quantification of Behavior,* Jackson Laboratory, ME (JG), an NIMH R01 MH132822 (MZ), a Klingenstein-Simons Early Investigator Award (MZ), a Whitehall Fellowship (MZ), a Sloan Fellowship (MZ), and a McKnight Scholars Award (MZ). Author M.Z. is supported as a Howard Hughes Medical Institute (HHMI) Freeman Hrabowski Scholar.

## CONFLICTS OF INTEREST

The authors declare no competing interests.

## STAR METHODS

### Key resources table

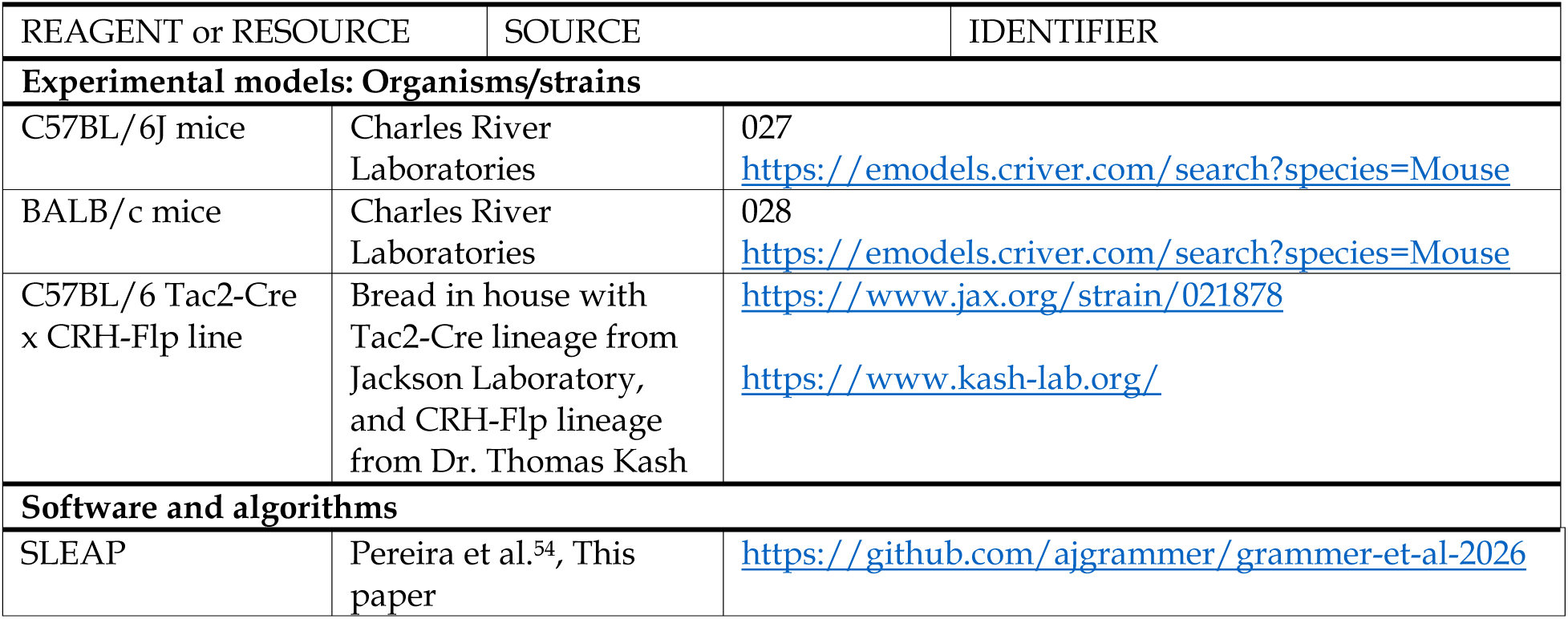
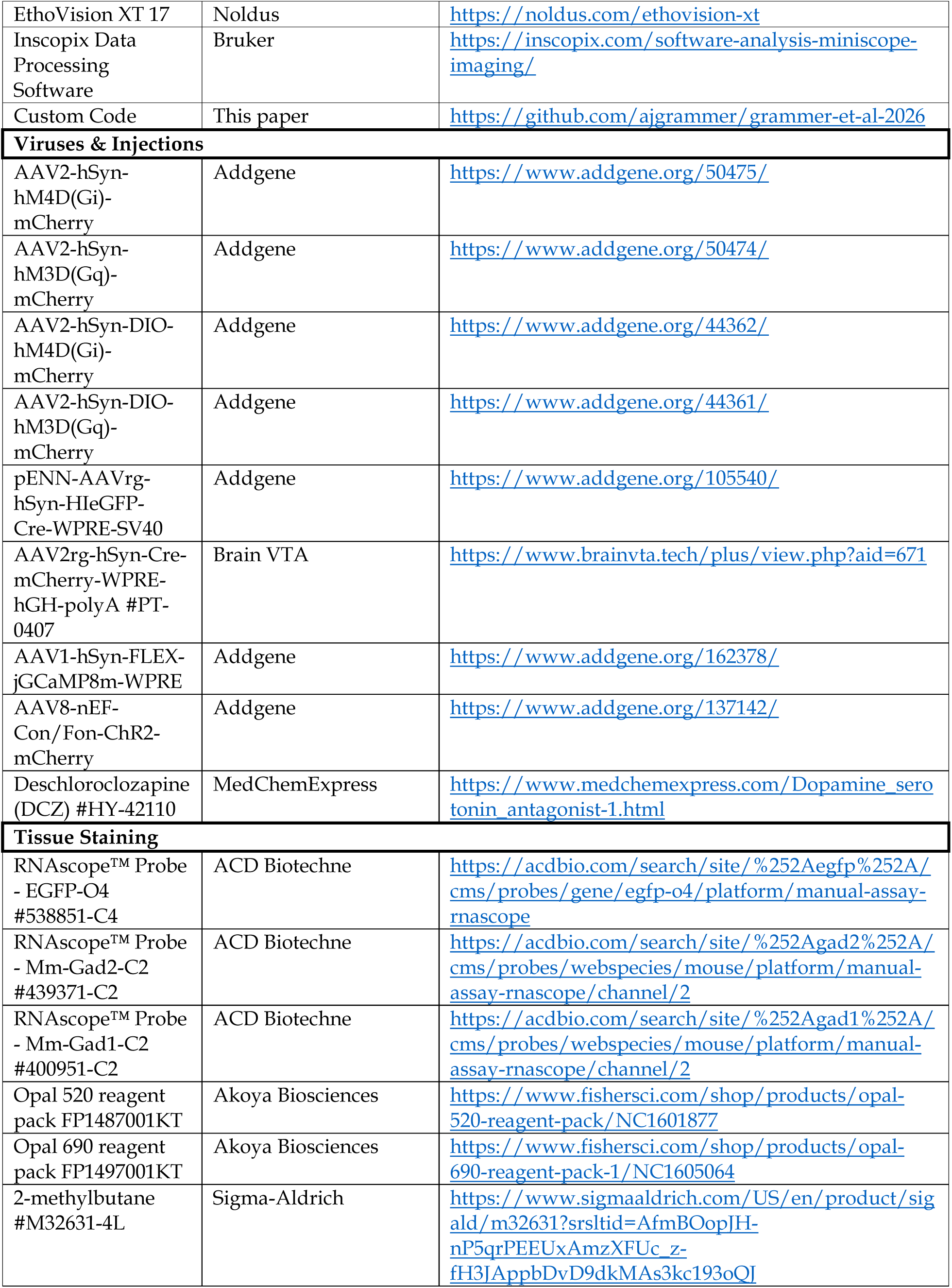
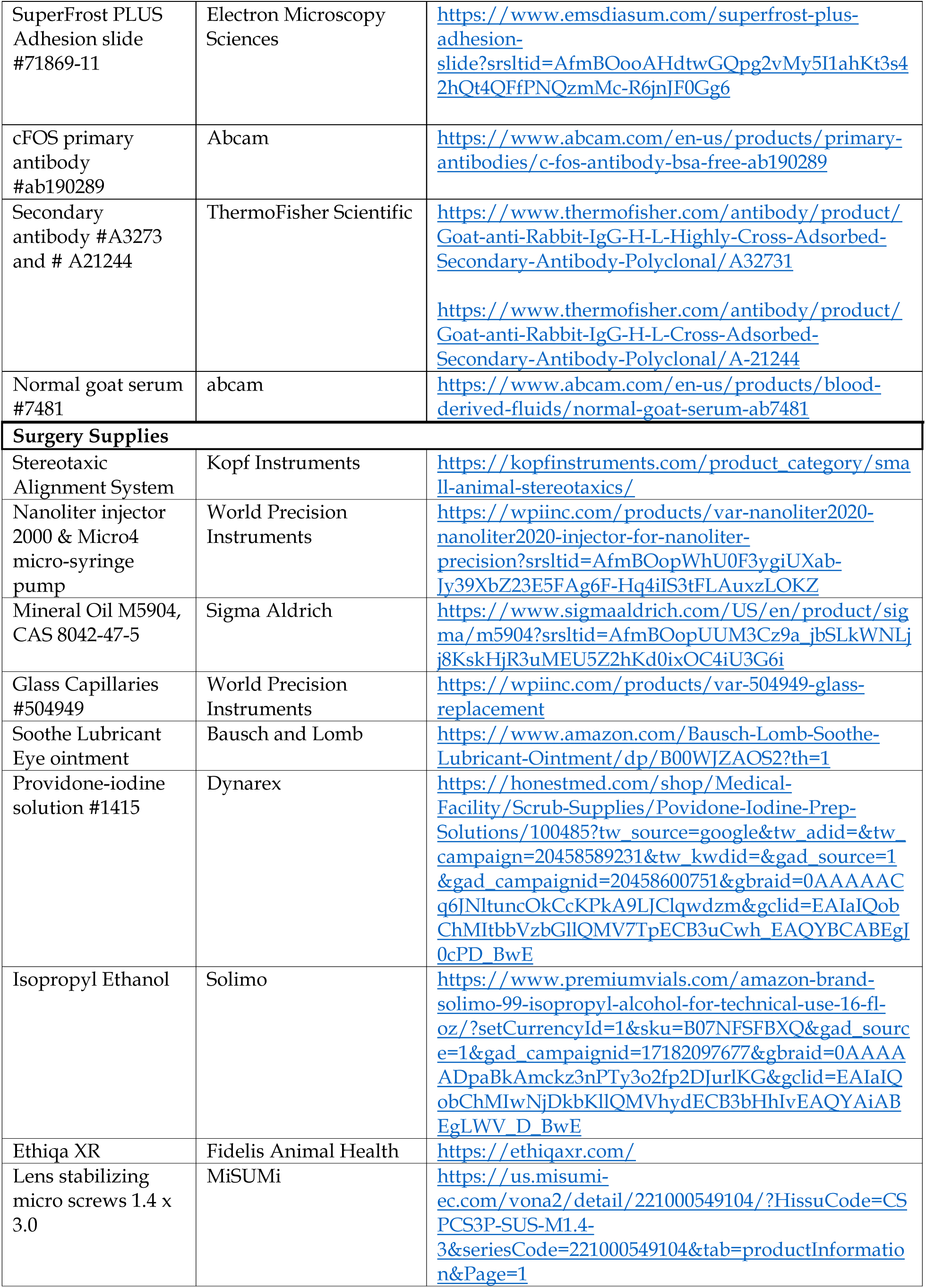
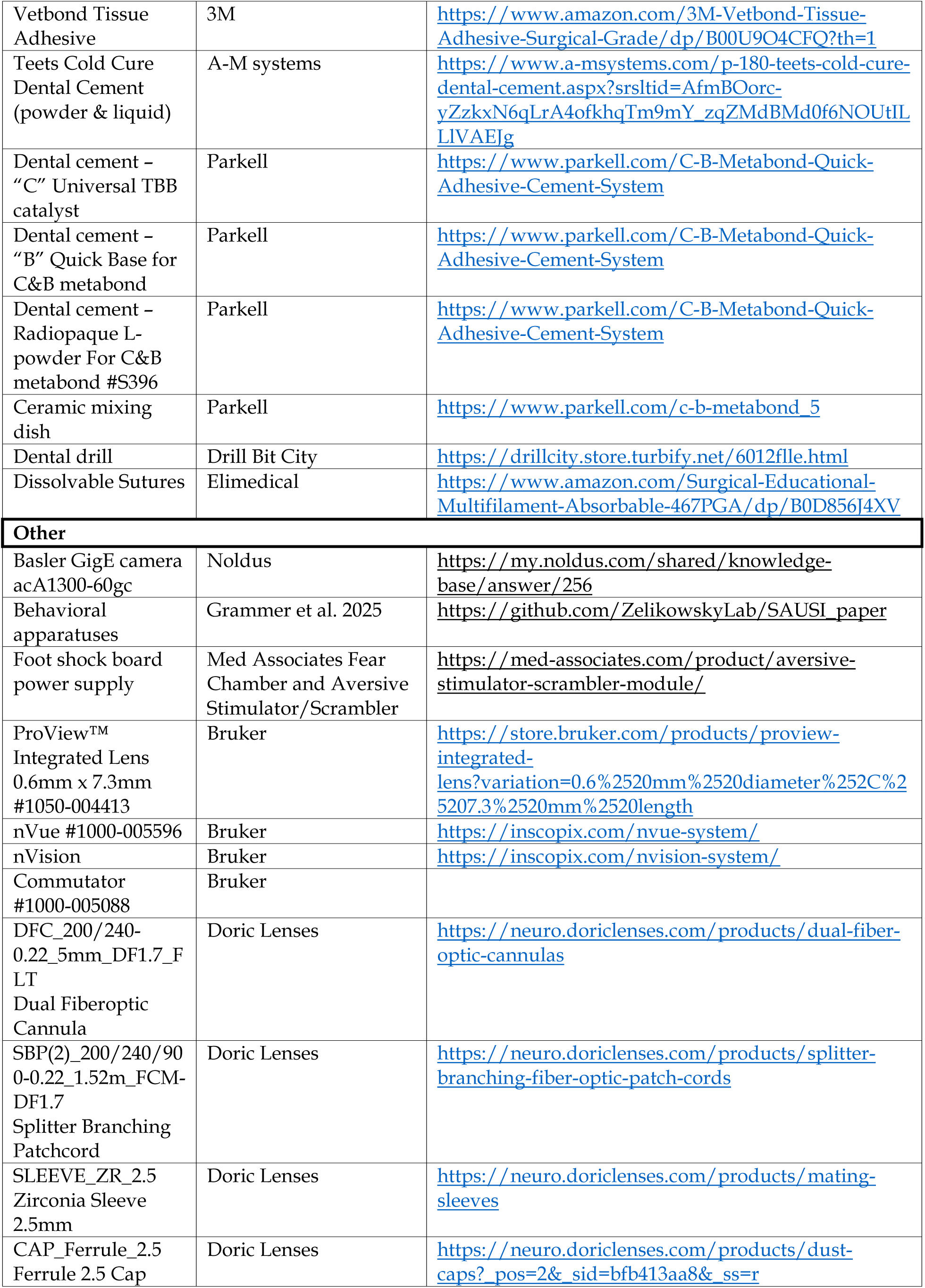
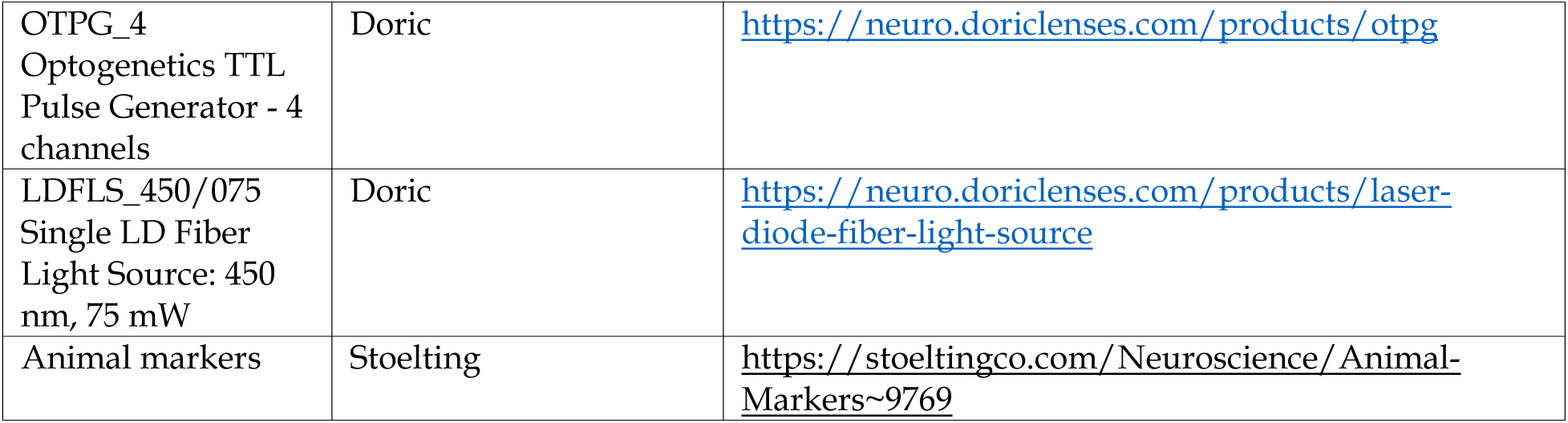

### Experimental model and study participant details

C57 BL/6 and BALB/c mice were housed in cages (Thoren type 9 https://thoren.com/?page_id=71) with Teklad pelleted bedding (https://www.inotivco.com/7084-pelleted-paper-contact-bedding) on a ventilated rack in a 12 hr light/dark cycle with unrestricted access to food (Teklad Global Soy Protein-Free Extruded Rodent Diet 2920X) and water. All animal work was reviewed for compliance and approved by the Institutional Animal Care and Use Committee of The University of Utah (IACUC protocol #00001504).

### Method details

#### Room conditions

A bar was hung above the SAUSI apparatus for mounting lamps, cameras, and commutators.

*Lighting*: 2 lamps facing upward, yellow hue, lux = 22-23 per chamber.

*Temperature*: 70-75 F in behavior suite.

*Handling*: plastic beakers (4” diameter, 5” tall) were used to transport mice from home cages into SAUSI. Tethered mice were briefly placed under anesthesia for attachment to cord.

On the first day of the SAUSI protocol, mice were weighed. Each experimental mouse was paired with a lower-weight BALB/c (2 weeks younger) that was sex matched. Each BALB/c was tested with both a group housed and isolated mouse in counterbalanced order. The hind region of BALB/c mice was colored with Stoelting animal markers to be identified throughout training and testing.

#### Mice

Wildtype mice were purchased from Charles River Laboratories. The Tac2-Cre x CRH-Flp line was bread in house with Tac2-Cre lineage from Jackson Laboratories and CRH-Flp lineage from the lab of Dr. Thomas Kash, backcrossed to C57BL/6Ns. The strain of experimental mice was C57BL/6N. The strain of conspecific mice was BALB/c. Experimental mice arrived at the University of Utah vivarium facilities at 5-6 weeks old. All mice were housed on a ventilated rack with dark, opaque plastic dividers stationed in between cages. These features were present to reduce the visual, auditory, and olfactory stimuli of surrounding cages for isolation. Isolated mice were singly housed starting at 6 weeks old and group housed mice were housed 3-4 to a cage. Only female mice were used throughout (except for figure S3) to recapitulate a social aversion phenotype across all three categories (social motivation, fear, hesitancy)^8^.

#### SAUSI protocol

SAUSI behavior was carried out in accordance with Grammer et al. 2025^8^.

The SAUSI apparatus contains two outer rooms (“chambers”), one empty (“home”) and one containing a conspecific (“social”). These rooms are connected by a narrow, transparent tunnel with a miniature footshock board on either side, which deter conspecifics from crossing over during pre-training and allows for observable decision making in the experimental mice.

To habituate mice to the apparatus, tunnel, and handling, experimental mice are placed in the arena for 5 minutes each on days 1 and 2. During this assisted habituation, experimenters gently guided the mouse through the tunnel every 30 seconds. On the third day, mice were placed in the apparatus and monitored to ensure that the mice can use the tunnel at will.

The conspecifics underwent a 5-minute 0.3mA foot shock deterrent session on the day prior to SAUSI testing to prevent them from crossing to the other side and blocking the tunnel entryway. During the deterrent session they received a 0.3mA foot shock any time they steppped on the board. In previous work, we found that this shock amplitude was sufficient to generate deterrence without changing the behavior of the conspecific or target mouse^8^. Tape was placed over the tunnel openings (or the tunnel block was used for the tether-compatible tunnel) during deterrent so that mice could not shelter in the tunnel during the first shock. Conspecifics were then used for testing (2 or more times) without additional deterrent training for 2+ weeks.

Our SAUSI social behavior test consisted of a 3-minute habituation period (where no other mice or stimuli were present in the chamber) followed by a 10-minute test phase with the conspecific present. Mice started on the same side during baseline and test phases, and the social chamber was always opposite to the start side (start side counterbalanced between mice). To transition between baseline and test phase, mice were briefly placed in their starting chamber with the tunnel blocked by a heavy object while the conspecific is placed in the opposite chamber. The test phase began once the tunnel block was removed. In the event that a mouse never entered the social chamber or did so for a limited amount of time (<10s), mice were placed in the social chamber for 10 additional minutes with the tunnel blocked to obtain social behavior data.

#### Resident Intruder and Optogenetics

To activate the oval nucleus of the BNST, Tac2-Cre x CRH-Flp c57BL/6 mice were bilaterally injected with 200nL (per side) of AAV8-nEF-Con/Fon-ChR2-mCherry virus at coordinates AP +0.5 ML +-0.85 DV -4.1 relative to best fit bregma^73^, just anterior to the cross section of coronal and sagittal skull sutures. A fiber optic cannula was also implanted in the same surgery at 0.1mm above the viral injection site.

Videos were recorded from an overhead camera under infrared light (43 cm from the cage floor) using MediaRecorder and manually scored in EthoVision. A box surrounded each cage during testing and contained a ledge to prevent mice from exiting the cage during testing^8^.

Mice were briefly placed under anesthesia to connect the fiber-optic patch cord to the implanted cannula. The RI tests began with a 3-minute baseline phase, where the experimental mouse was alone in the home cage. This was followed by a 3-minute social baseline phase with laser off, then a 12-minute test phase with alternating light on/off for 3-minute segments (counterbalanced). For stimulation, 5ms pulse trains at 20 Hz, 450 nm light, and 5 mW at fiber tip were used.

Cages used (Thoren type 9- https://thoren.com/?page_id=71) were 19.56 x 30.91 x 13.34 (cm.) with a floor area of 435.7 (Sq. cm.). Bedding that mice were continuously housed in (Teklad pelleted bedding - https://www.inotivco.com/7084-pelleted-paper-contact-bedding). Nestlets were removed for visibility throughout the task. Food and water were not accessible during the task.

Group-housed mice that were not actively being tested were placed together in a clean holding cage. Each group-housed mouse shared an “untested” holding cage with their cage mates prior to testing. After being tested, they were placed in another clean “tested” holding cage so that no untested mice were exposed to tested mice.

All mice were group housed (male n=4; female n=5) for resident intruder optogenetic experiments.

#### Surgeries

Mice were placed under Isoflurane anesthesia at an induction rate of 4-5% and a maintenance rate of 1-2%. Once mounted on the stereotaxic surgery rig, a 3.25mg/kg subcutaneous injection of Ethiqa XR was given for analgesia. Eye lubrication was placed over the eyes. Hair was removed using Nair and the skin was sterilized using 3x providone-iodine solution and 70% isopropyl alcohol. A small incision was made on the scalp, and the skull was levelled. Craniotomies at the viral injection coordinates were created using a dental drill. A pulled glass capillary (∼50 micron diameter tip) was filled with mineral oil and virus was pulled into and expressed from the capillary using a nanoliter injector and micro syringe pump. All viruses were injected at a rate of 50nL/min and the capillary remained in the brain for 1 minute after each injection before quick removal. For surgeries without implants, incisions were closed with dissolvable sutures. For optogenetic and microendoscope surgeries, the skull was additionally primed by scratching the surface with the back of a scalpel blade, using Vetbond to seal the edges of the skin, and screwing 2 micro screws into the skull on opposing sides to the fiber optic cannula (or GRIN Lens) implant. Implants were stabilized with dental cement. Mice were given at least 3 weeks for viral expression and recovery time.

#### DREADDs experiments

Female C57BL/6N mice were either group housed or isolated at 6 weeks old. Viral injection surgeries were performed at 6-7 weeks old. To perturb the pan-neuronal adBNST population, mice were bilaterally infused (150nL per side) with an inhibitory DREADDs virus (AAV2-hSyn-hM4D(Gi)-mCherry) (Fig. 2) or an excitatory DREADDs virus (AAV2-hSyn-hM3D(Gq)-mCherry) (Fig. S2) at coordinates AP +0.95 ML +-0.85 DV -4.1. To silence the adBNST to NAc projections specifically (Fig. 3), a Cre-expressing retrograde virus (pENN-AAVrg-hSyn-HI-eGFP-Cre-WPRE-SV40) was infused bilaterally (200nL per side) into the NAc (AP +1.98 ML +-0.75 DV -4.5) and a Cre-dependent DREADDs virus (AAV2-hSyn-DIO-hM4D(Gi)-mCherry) was infused bilaterally (150nL per side) into the adBNST (AP +0.95 ML +-0.85 DV -4.1) in the same surgery^74^. All DREADDs coordinates were relative to traditional bregma^73^, at the intersection of coronal and sagittal skull sutures. At 10 weeks old, mice underwent SAUSI behavioral testing. SAUSI testing, as described above, with 3 days of habituation followed by the first social test. Then after one day of rest, the same mice are retested in the SAUSI assay for a within subjects design. On the day of the social test, mice receive an intraperitoneal (IP) injection of either 0.25mg/kg Deschloroclozapine (DCZ) or vehicle (DMSO + saline) 10 minutes prior to the start of the baseline phase. Mice receive the opposite injection on the second social test day (order counterbalanced). After behavior, brains were extracted, post-fixed in 4% paraformaldehyde, sliced in 40-micron sections, dry mounted onto slides, stained with DAPI and cover-slipped. A Leica DM2000 fluorescent microscope was used to take 10x images for viral placement confirmation in the NAc and BNST.

We also tested whether the 0.25mg/kg DCZ dose had any off-target effects in viral control experiments (Figure S2A), and found none (Figure S2B-D). A higher DCZ dose of 1mg/kg (Figure S2E) did have off- target effects as indexed by reduction in social chamber preference (Figure S2F), a trend for increased social freezing (Figure S2G), but no change in social hesitancy (Figure S2H). For testing off-target effects, the same design as above was used, but mice did not undergo surgery for viral injection.

#### Fluorescent in situ hybridization

Mice were group housed or isolated for 4 weeks starting at 6 weeks old. Mice were placed under isoflurane anesthesia prior to rapid decapitation. Brians were placed in 2-methylbutane for ∼10 seconds and then were placed in histology molds covered in tin foil at -80°C. Brains were sliced into 20-micron sections using a cryostat and dry mounted onto SuperFrost PLUS slides. Each slide contained 1 slice (2 hemispheres) per mouse and 4 mice total. Slide boxes were wrapped in Seran wrap and placed in a Ziploc bag for airtight storage at -80°C. The RNAscope assay was performed according to Advanced Cell Diagnostics protocol^75^. Slides were imaged with a Keyence BX-X800 using a 10x objective. See quantification details below.

Representative images were taken at 10x and 40x on an Echo Revolution Confocal Microscope.

#### DREADDs Verification

We confirmed that the Designer Receptor Exclusively Activated by Designer Drugs (DREADDs) approach with a 0.25mg/kg dose of Deschloroclozapine (DCZ)^76^ was effective at silencing or activating neurons in the adBNST using cFOS as an indirect marker for neural activity^77^ (Figure S2M-O). Mice were injected with either DCZ or vehicle and returned back to the home cage. Mice were then placed under 5% anesthesia and sacrificed with rapid decapitation 105 minutes later, and brains were post-fixed in 4% paraformaldehyde.

This 105 minute time window was chosen to accommodate peak cFOS expression (90 minutes)^78^ + 15 minutes to simulate the SAUSI behavior timeline (where mice are injected with DCZ or vehicle 10 minute prior to the start of the baseline phase, which is ∼15minutes prior to the test phase). Following dehydration through 15% and 30% sucrose solutions, brains were embedded in Optimal Cutting Temperature (OCT) compound and stored at -80°C. Brains were then sliced at 40 microns thick as free floating sections in 1x Phosphate Buffered Saline (PBS) using a cryostat.

Free floating immunohistochemistry^79^ was performed to stain for cFOS expression. Rabbit anti cFOS abcam #190289 was used as a primary antibody. Slices were then blocked in normal goat serum. For brains with pan-neuronal viral expression (tagged with mCherry), goat anti-rabbit AlexaFluor 488 was used as a secondary. For brains with projection specific viral expression (tagged with both mCherry and eGFP), goat anti-rabbit AlexaFluor647 was used as a secondary. Images were taken with a Keyence BX-X800 using a 10x objective. See quantification details below.

#### Microendoscope experiments

Female mice were isolated and underwent stereotaxic surgery at 6 weeks old to implant a Bruker ProView™ Integrated GRIN Lens 0.6mm x 7.3mm with an integrated base plate over the Bed Nucleus of the Stria Terminalis (lens lowered to AP +0.95, ML +- 0.85, DV -4.05). Mice were also injected unilaterally with a retrograde Cre-expressing virus in the Nucleus Accumbens (AAV2rg-hSyn-Cre-mCherry-WPRE-hGH-polyA; 200nL; AP +1.98 ML +-0.75 DV -4.5) and a Cre-dependent virus encoding the calcium indicator GCaMP8m (AAV1-hSyn-FLEX-jGCaMP8m-WPRE; 300nL; AP +0.95 ML +-0.85 DV -4.1) in the same surgery. All microendoscope coordinates were relative to traditional bregma^73^, at the intersection of coronal and sagittal skull sutures. Mice were given approximately 6 weeks to recover before behavior experiments.

For behavioral testing, the tether-compatible open-top tunnel was used for SAUSI^8^. Mice were briefly placed under isoflurane anesthesia and attached to a miniature microendoscope (Bruker nVue) which was wired to a commutator connected to a Data Acquisition System (DAQ box). Synced videos were captured using the nVision system.

All mice were habituated with the tether throughout the 3 SAUSI habituation days. All SAUSI protocols were identical to previous testing except mice were handled by tail rather than in transport beakers and the experimenter was present in the room throughout each session to ensure there were no problems with the cord. After behavior, brains were extracted, post-fixed in 4% paraformaldehyde, sliced in 40-micron sections, dry mounted onto slides, stained with DAPI and cover slipped. A Leica DM2000 fluorescent microscope was used to take 10x images for viral placement confirmation in the NAc and BNST. All viral confirmations were performed by a blinded experimenter.

Behavior videos were converted to .mp4 files using the Inscopix Data Processing Software. Videos were tracked and hand scored using EthoVision XT 17. Data was graphed and analyzed using custom code https://github.com/ajgrammer/grammer-et-al-2026 and GraphPad Prism.

### Quantification and statistical analysis

Data were presented as mean ± Standard Error of Mean (SEM) with individual values displayed. At least 2 experimental replications were conducted in each main figure.

#### Behavior Analysis

| **Behavior** | **Event Type** | **Description** | **equation** |
| --- | --- | --- | --- |
| Sniffing | Start-stop | Sniffing the conspecific |  |
| Conspecific sniff* | Point event | Any new sniff bout by the conspecific |  |
| First Tunnel Sheltering | Start-stop | In tunnel before first time entering the social chamber (can be multiple times before entering). must have all 4 paws in tunnel. For microendoscope experiments: stopping in the tunnel. |  |
| Reverse in Tunnel | Start-stop | Backing up in the tunnel after all 4 paws entered. |  |
| Tunnel Approach | Start-stop | In the tunnel approaching the social chamber. Must have all 4 paws in tunnel. |  |
| Tunnel Retreat | Start-stop | In the tunnel approaching the home chamber. Must have all 4 paws in tunnel. |  |
| Social Freeze | Start-stop | freezing in response to being sniffed by the conspecific. Can be during or just after being sniffed by the conspecific (within 1 second). Can include shifting body away from conspecific with a frozen posture | $\frac{\text{social freeze bouts}}{\text{conspecific sniff}} * 100\%$ |
| Social Reactivity | Point event | experimental mouse reacts when conspecific sniffs it (within 1 second of the end of a sniffing bout from the conspecific). Reacting can include retreating, darting, vigilance, flinching, or jumping. | $\frac{\text{social reactivity bouts}}{\text{conspecific sniff}} * 100\%$ |
| Prosocial Initiation | Point event | The experimental mouse approaches the conspecific to sniff and closes the gap between them/ comes from more than one body length distance away. The conspecific does not close the gap. Marked at the point of arrival. | $\frac{\# \text{ initiations}}{\text{social chamber time (s)}}$ |
| Social Chamber Preference | Start-stop | Preference to spend time in the social chamber. | $\frac{\text{social chamber time (s)}}{\text{total time (s)}} * 100\%$ |
| freeze | Start-stop | Freezing not in response to being sniffed by the conspecific. (at least 0.5 seconds duration) |  |
| aggression | Start-stop | Attack, biting, mounting |  |

*\*all behaviors are referencing the experimental mouse except for conspecific sniff.*

Videos were recorded in MediaRecorder or the Bruker nVision platform. The top view perspective was used for automated and manually scored behavior with EthoVision^80^ as well as postural data tracking in SLEAP^54^.

#### Postural Data Collection and Analysis

SLEAP ^54^ was the program used for postural tracking. All videos used for tracking were from the top camera view and were trimmed to only include the test phase. A 10-point skeleton was used for each the experimental and conspecific mouse including nose, 2x ears, skull base, 2x arms, mid-back, 2x legs, tail base. 190 frames were manually labelled for training. All training, tracking, and post-track editing were performed in the GUI. Bottom-Up processing was used for training with default settings. For inference, tracker was set to “flow”, clean_instance_count was set to 2, track_window was set to 30, and all remaining settings were left as default. Each SLEAP tracked video was run through a custom post-correction python workflow (https://github.com/ajgrammer/grammer-et-al-2026), which added a layer of identity track using mouse size, color and motion to correct long-distance travel, identity swaps, & double labelling of a single mouse.

After this iterative pipeline was used on predicted videos, tracking was manually edited in the SLEAP GUI to ensure that identities were correct throughout.

Python was used to generate all SLEAP downstream analysis (code available here https://github.com/ajgrammer/grammer-et-al-2026). 20+ features (ex: nose-nose distance, approach velocities, body lengths, facing angles, etc.) were calculated from the raw SLEAP output and were used to generate low dimensional embedding. Features were z-scored and embedded into two dimensions using Uniform Manifold Approximation and Projection (UMAP; *n_neighbors* = 50, *min_dist* = 0.1, Euclidean distance). Figure 1 (G-T) showed “social” UMAPs and subsequent analysis, where only frames where mice were <2 body lengths distance apart were used. Figure S1 used data from all frames (social and non-social) from the SAUSI test to generate UMAPs and subsequent analyses. Figure 1 facing angle analysis (U-V) used all frames (social and non-social). A UMAP occupancy histogram was smoothed with a Gaussian filter (σ = 2 grid units), and local density maxima were identified using a minimum peak separation of 10 grid units. Watershed segmentation was then applied to the inverted density map to define separate regions.

To quantify the spatial distribution of manually annotated behaviors overlayed on the UMAP, the root-mean-square (RMS) was taken from the distances of all points from their centroid (arithmetic mean of all points). These distances were averaged per mouse, accounting for any difference in the total number of behavior points present in the UMAP.

To decode housing condition from unsupervised clustering, a logistic regression model was used, with watershed region occupancy as input. Models were fit using L2 regularization (liblinear solver), median imputation of missing values, region occupancy standardization, and balanced class weights. Performance was evaluated across 20x repeated stratified (ie randomized but balanced between housing condition) train/test splits (80/20). The receiver operating characteristic area under the curve (ROC-AUC) is the average from each iteration of the model. Region 1 was excluded from interpretation due to the small quantity of frames (Figure 1T).

We used facing angle data to quantify the relationship between social orientation and behavior. Facing angle was calculated as the absolute value of 0 to Pi (radians), where 0 indicates the other mouse is directly in front of the mouse of interest, and Pi is directly behind. These values were then binned into 3 categories (front, side, behind). Probability of each behavior occurring (# behavior-positive frames in angle bin / # total frames in angle bin) was plotted as a function of facing angle for both the experimental and conspecific mouse. The area under the curve (AUC; trapezoidal integration) analysis was computed per facing angle bin. Group differences were assessed with a 2-factor ANOVA (housing condition x facing angle bin) with Holm post-hoc correction.

Watershed region descriptions were deciphered using information from image frame examples from regions, SLEAP features, and manual annotation overlays on the UMAP.

GraphPad PRISM was also used to generate graphs and statistics.

#### Cell Counting

FIJI^81^ was used for all microscopy image quantification. Briefly, regions of interest were drawn around the entire dBNST, the anterodorsal subregion (including anteromedial, anterolateral, and juxtacapsular nucleus)^28^, and the oval nucleus. Expression thresholds were set for each channel. Then masks were made using 25-infinity pixels and 0.5-1 circularity. These masks were used for automated cell counting. Blind experimenters also oversaw this analysis and adjusted with hand counting if the masks were not representative of true cell count (for example due to low expression in the image).

#### In vivo Calcium Imaging

Videos were processed in Inscopix Data Processing Software including down sampling, spatial filtering, motion correction, and automated cell detection with Constrained Non-negative Matrix Factorization for Endoscope (CNMF-E). Delta F/F activity traces were exported as csv file and temporally aligned to behavior data. Calcium activity traces were then normalized using a rolling baseline correction, median filtering, light temporal smoothing (Savitzky-Golay) and z-scoring. Peri-event time histograms (PETHs) were created for each behavior with a 10 second window (5 second pre, 5 second post). For a neuron to be classified as tuned, it must have met the following 2 criteria: 1) its peak response post onset must have exceeded 3 times the baseline mean, 2) reached statistical significance when compared to a shuffled control. Statistical significance was assessed using a nonparametric percentile test, where activity must have exceeded the 97.5^th^ percentile of the shuffled distribution (one-tailed, alpha = 0.025). A circular-shift shuffle procedure was repeated 200 times to create the null shuffled distribution. For each neuron, the calcium trace was circularly rotated by a random offset while preserving temporal structure, and peri-event activity was recomputed using the original behavioral timestamps. All data shown are from cells classified as tuned and excited by behavior. Significant activation post behavior onset in PETH graphs was performed by comparing the average signal (0 to 5 second window) per neuron to its shuffled control.

For population recruitment calculations, the fraction of total neurons that exceeded the baseline (pre-onset) mean was graphed for each frame of the PETH window. Because this was a binary metric (exceeded baseline or did not exceed baseline), the pre-onset window had an average value of 0.5 (chance).

For our predictive decoding data, a lagged logistic regression model was used with calcium traces as the input and each binary behavior (present 1, absent 0) as a separate output. For the lag, activity from each neuron was concatenated across the current frame and the previous 30 frames (31 total time points), generating a feature matrix that captured recent temporal dynamics of the neural population. Balanced class weighting was incorporated to compensate for unequal frequencies of behavioral and non-behavioral frames. Neural features were standardized using z-score normalization. Models were trained using the LIBLINEAR solver. Cross-validation was achieved using a 5-fold expanding-window time-series method.

This method was chosen for rigor - to avoid predicting on frames quazi-identical to test frames due to the nature of highly correlated neural activity and behavioral data. In each fold, the classifier was trained on sequential observations preceding the test interval and evaluated on a subsequent, non-overlapping test block. Predictions from each iteration were concatenated to generate a single prediction vector for each behavior and mouse. Decoder performance was quantified using receiver operating characteristic area under the curve (ROC-AUC). To estimate chance-level decoding performance, shuffle-control analyses were performed independently for each behavior. Behavioral labels were randomly permuted while preserving the neural activity matrix, and the complete decoding procedure was repeated for each shuffle iteration.

Shuffle performance was quantified using ROC-AUC, generating a null distribution for each mouse and behavior. Real vs. shuffled ROC-AUC comparisons were assessed using paired t-tests. Multiple comparisons across behaviors were corrected using the Benjamini–Hochberg false discovery rate (FDR) procedure^82^.

#### Exclusion Criteria

Hesitancy scores were excluded if the conspecific stepped on the footshock board (and received a shock) during the first tunnel entrance of the experimental mouse, which can impact latency to approach, first tunnel sheltering, and reversing in tunnel. If conspecific aggression occurred, then behavior data only prior to the first bout of aggression was included. If a conspecific entered the tunnel at any point during testing, that trial was excluded. Group-housed cages were excluded if there was overt aggression in the home cage. Spatially isolated watershed regions were excluded due to low frame count (<5 frames). For DREADDs experiments, mice were excluded if there was viral spread into the oval nucleus of the BNST in at least one hemisphere, or an inaccurate targeting of the retrograde virus in the nucleus accumbens. For microendoscope experiments, mice were excluded if any virus or lens placement was not accurate to the targeted region, or if no valid cell activity was recorded.

## Notes

https://github.com/ajgrammer/grammer-et-al-2026

### Competing Interest Statement

The authors have declared no competing interest.

