## supplemental figures for "Extended amygdala orchestrates social motivation in socially isolated mice"

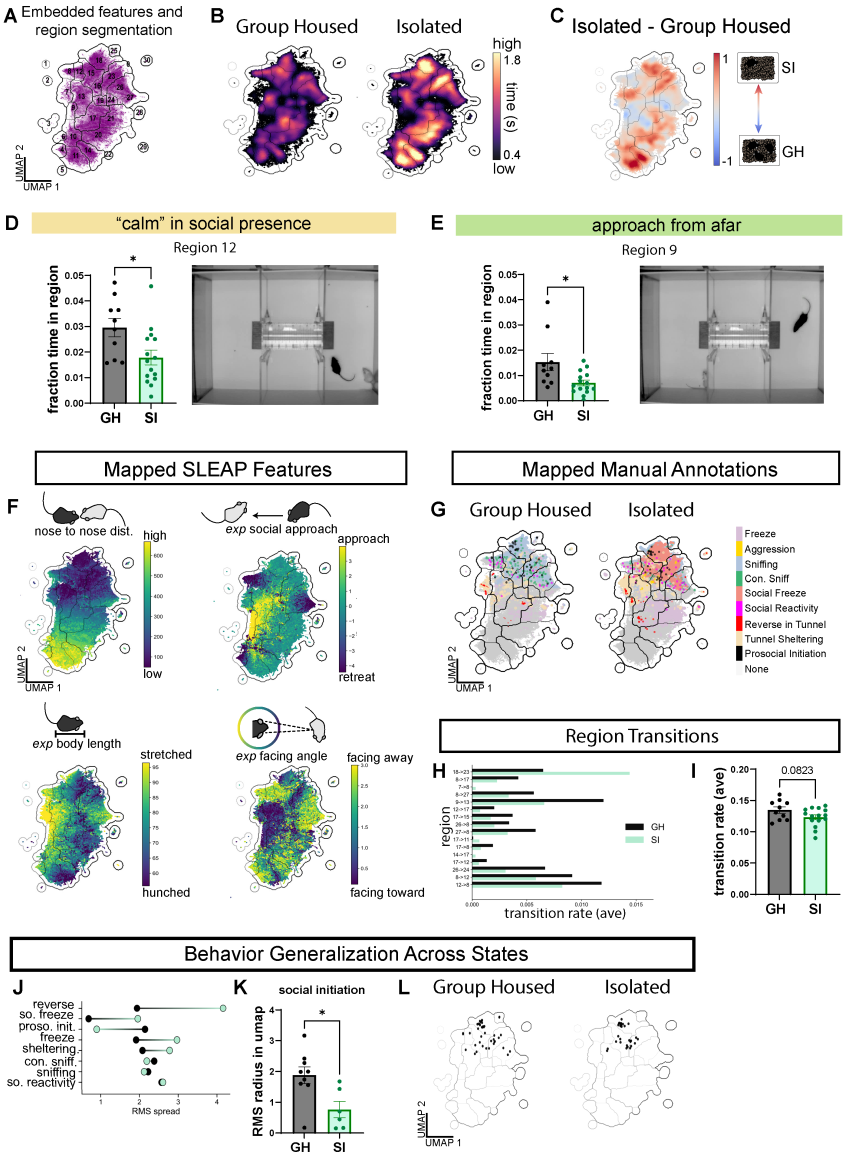


**Figure S1. | Behavior quantification with social and non-social frames.** (**A**) Entire assay (including all frames) UMAP and segmentation from all mice. (**B**) Time occupancy in UMAP regions by housing condition. (**C**) Change in UMAP occupancy between housing conditions. (**D**) SI mice spend less time occupying region 12 than GH mice (Welch's t-test, $p<.05$) (left); still frame from behavior video for region 12 (right). (**E**) SI mice spend less time occupying region 9 than GH mice (Welch's t-test, $p<.05$) (left); still frame from behavior video for region 9 (right). (**F**) Calculated SLEAP features embedded in UMAP. (**G**) Manual annotations embedded in UMAP. (**H**) Average transition rate between regions (of the top 16 transitions) by housing condition. (**I**) Across all frames, socially isolated mice tend to transition between regions less often than group housed mice (Welch's t-test, $p=.08$). (**J**) Averaged distribution of each behavior across UMAP by housing condition. (**K**) Socially isolated mice show tighter distribution of prosocial initiation compared to group housed mice (Welch's t-test, $p<.05$). (**L**) Visualization of prosocial initiation UMAP distribution differences across housing conditions. *n=10 group housed, n=15 isolated for all data in Figure S1. All error bars represent mean +/- standard error of the mean. GH: group housed; SI: socially isolated;*

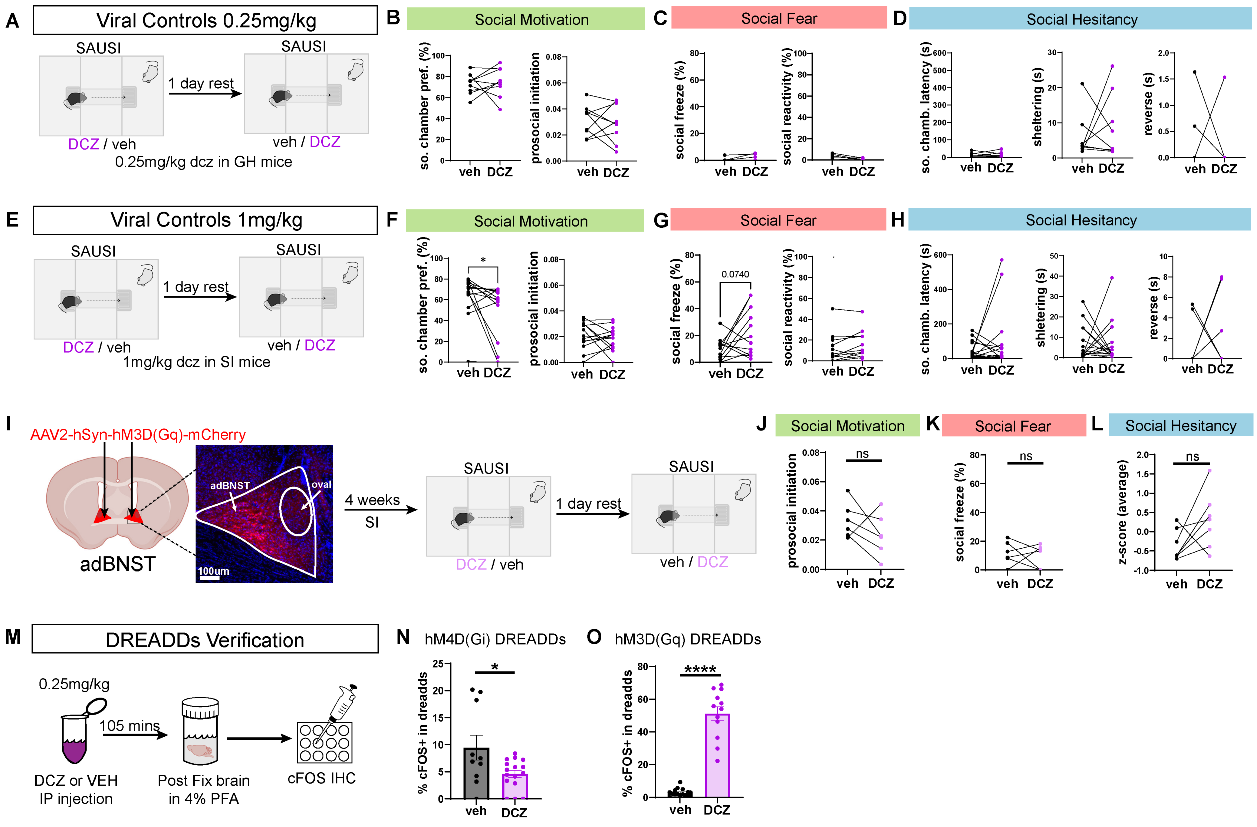


**Figure S2. | DREADDs with 0.25mg/kg DCZ is effective with no off-target effects.** (**A**) Experimental design for DCZ control experiment with 0.25mg/kg DCZ dose. (**B-D**) 0.25mg/kg injection of DCZ in wildtype mice has no off-target effects on behavior. (**E**) Experimental design for DCZ control experiment with 1mg/kg DCZ dose in wildtype mice. (**F**) 1mg/kg DCZ dose may have an off-target effect on social motivation as indexed by reduced social chamber preference (n=15; paired t-test, p=.04). (**G**) 1mg/kg DCZ dose trends towards an increase in social fear as indexed by social freezing (n=12; paired t-test, p=.07). (**H**) 1mg/kg DCZ dose has no effect on social hesitancy behavior. (**I**) Experimental design for activating adBNST during behavior & viral targeting of adBNST. (**J**) Activating adBNST in SI mice has no effect on prosocial initiation (n=6; paired t-test, p=.25), (**K**) social fear (n=6, p=.75) or (**L**) hesitancy (n=7, p=.12). (**M**) Experimental design for testing the efficacy of DREADDs manipulations. (**N**) for inhibitory DREADDs, cFOS expression is significantly reduced when DCZ is administered compared to vehicle control (n=6 mice vehicle, n=8 mice DCZ; two hemisphere replicates per mouse; Nested independent samples t-test, p=.038). (**O**) For excitatory DREADDs, cFOS expression is significantly increased when DCZ is administered compared to vehicle control (n=8 mice vehicle, n=6 mice DCZ; two hemisphere replicates per mouse; Nested independent samples t-test, p<.0001). *All error bars represent mean +/- standard error of the mean. GH: group housed; SI: socially isolated; DCZ: deschloroclozapine; veh: vehicle.*

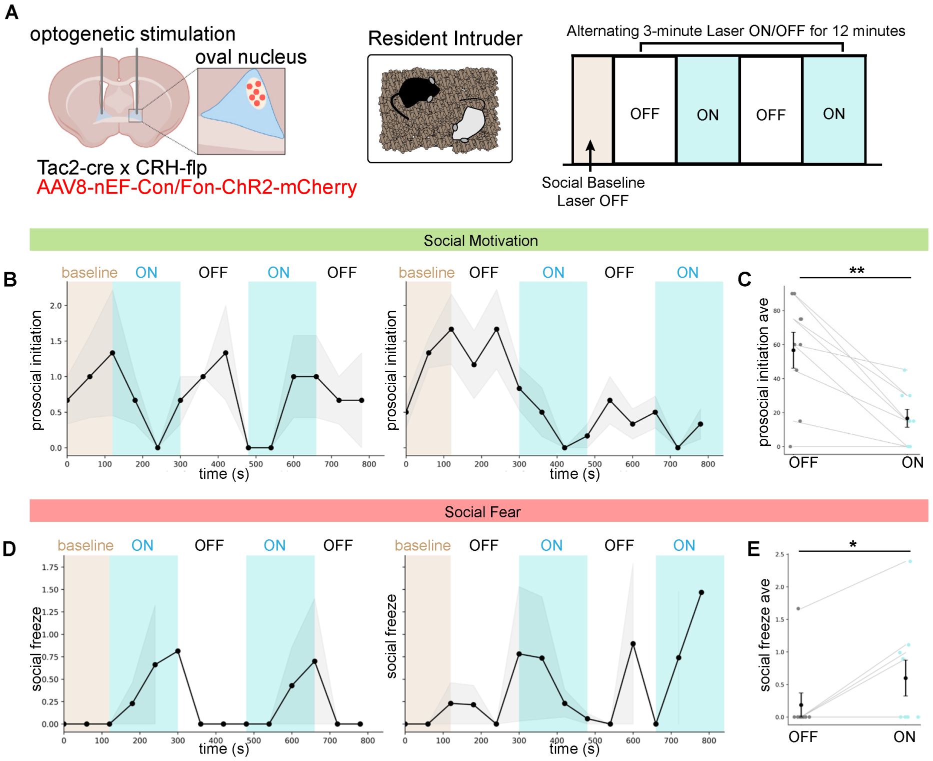


**Figure S3. | dBNST oval nucleus activation exacerbates social aversion.** (**A**) Experimental design for optogenetic activation of the oval nucleus. (**B**) Prosocial initiation is reduced by light-on stimulation of the oval nucleus, shown over time (left) and bins averaged (minutes 2-3 to exclude transition between ON/OFF bins) (right) (p=.002). (**C**) Social freezing (time/#conspecific sniffs) is increased by light-on stimulation of the oval nucleus, shown over time (left) and bins averaged (minutes 2-3 to exclude transition between ON/OFF bins) (right) (p=.038). *n=9, paired t-test. All error bars represent mean +/- standard error of the mean.*

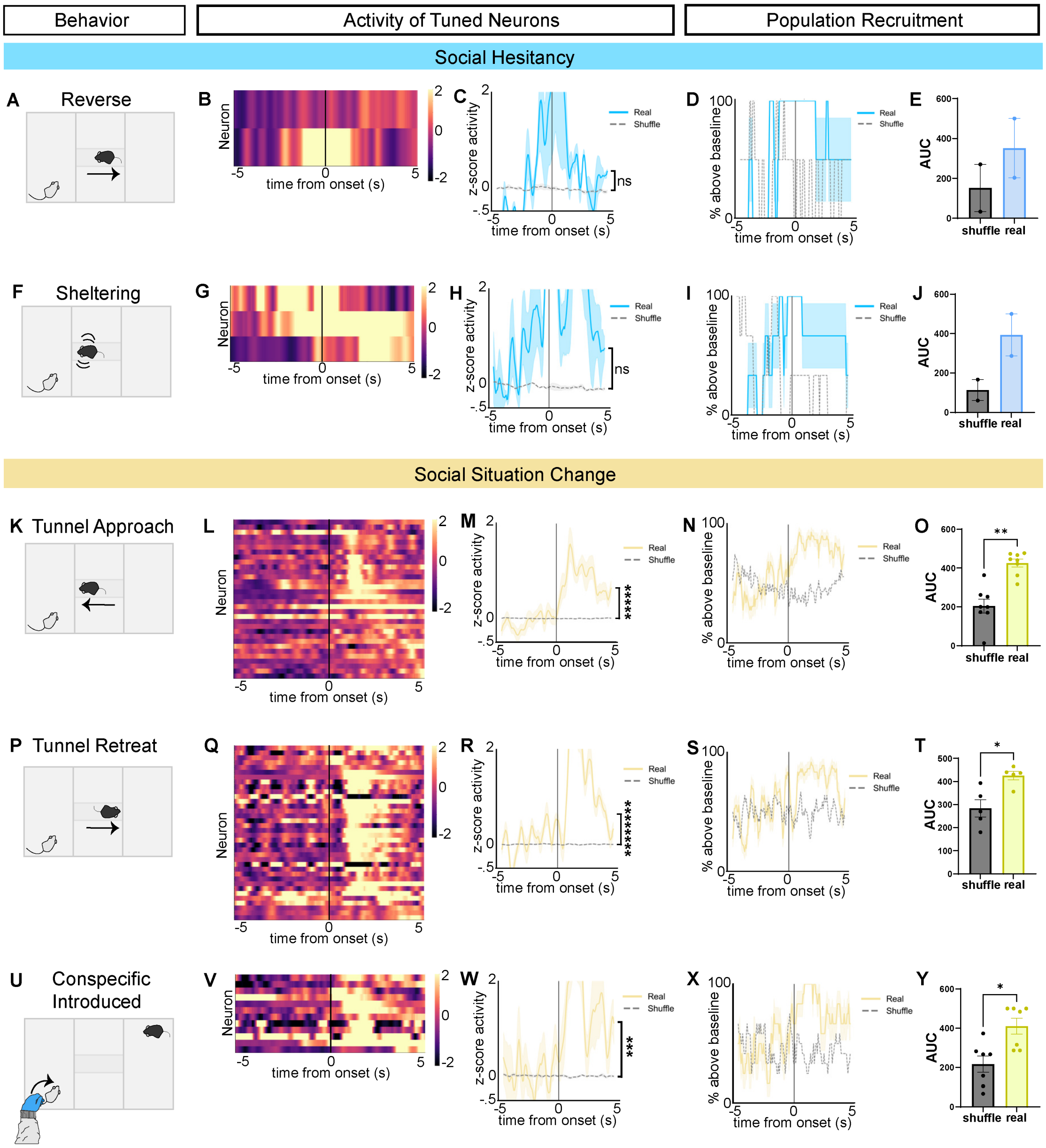


**Figure S4.** | **adBNST→NAc ensembles respond to social context changes.** (**A**) Behavior (reverse in tunnel) corresponding to neural activity analysis in panels B-E. (**B**) Heatmap of neurons tuned to reversing in the tunnel. (**C**) Neural activity PETH showing neurons tuned to reversing in tunnel, not significantly different from shuffled control (n=2 neurons; Wilcoxon signed-rank test, p=0.5). (**D**) Population recruitment PETH for reverse in tunnel. (**E**) No difference between population recruitment for reverse in tunnel between real and shuffled control data (n=2 mice; paired t-test, p=.09). (**F**) Behavior (sheltering) corresponding to neural activity analysis in panels G-J. (**G**) Heatmap of neurons tuned to sheltering. (**H**) Neural activity PETH showing neurons tuned to sheltering, not significantly different from shuffled control (n=3 neurons; Wilcoxon signed-rank test, p=0.25). (**I**) Population recruitment PETH for sheltering. (**J**) No difference between population recruitment for sheltering between real and shuffled control data (n=2 mice; paired t-test, p=.12). (**K**) Behavior (tunnel approach) corresponding to neural activity analysis in panels L-O. (**L**) Heatmap of neurons tuned to tunnel approach. (**M**) Neural activity PETH showing neurons tuned to tunnel approach, significantly increased compared to shuffled controls (n=32 neurons; Wilcoxon signed-rank test, p<.00001). (**N**) Population recruitment PETH for tunnel approach. (**O**) Coordinated response from population tuned to tunnel approach significantly differed from shuffled control data (n=8 mice; paired t-test, p=.003). (**P**) Behavior (tunnel retreat) corresponding to neural activity analysis in panels Q-T. (**Q**) Heatmap of neurons tuned to tunnel retreat. (**R**) Neural activity PETH showing neurons tuned to tunnel retreat, significantly increased compared to shuffled controls (n=36 neurons; Wilcoxon signed-rank test, p=6.02e$^{-9}$). (**S**)

Population recruitment PETH for tunnel retreat. (**T**) Coordinated response from population tuned to tunnel retreat significantly differed from shuffled control data (n=5 mice; paired t-test, p=.025). (**U**) Behavior (Conspecific introduced to social chamber) corresponding to neural activity analysis in panels V-Y. (**V**) Heatmap of neurons tuned to Conspecific introduction. (**W**) Neural activity PETH showing neurons tuned to conspecific introduction, significantly increased compared to shuffled controls (n=13 neurons; Wilcoxon signed-rank test, p<.001). (**X**) Population recruitment PETH for conspecific introduction. (**Y**) Coordinated response from population tuned to conspecific introduction significantly differed from shuffled control data (n=7 mice; paired t-test, p=.016). *Individual data points represent values per mouse. All error bars represent mean +/- standard error of the mean. AUC: area under the curve.*
